# Insulin Modulates Food Valence Through Astrocytes

**DOI:** 10.64898/2026.11.10.765591

**Authors:** J Zegarra-Valdivia, E Fernandez de Sevilla, M.Z. Khan, J Pignatelli, I Torres Aleman

## Abstract

Brain cells involved in assigning affective value (i.e.: valence) to stimuli modulating brain states are not yet fully identified. Since astrocytes modulate brain states, and insulin regulates feeding behavior and exerts its central actions in part through astrocytes, we analyzed the role of the insulin receptor in astrocytes in assigning valence to food. We used the modulatory actions of food on anxiety and fear, as proxies of the affective influence of food on mood. Mice lacking insulin receptors (IR) in GFAP astrocytes (GFAP-IR KO mice) showed increased anxiety and fear in response to food, while control littermates showed the opposite behavior. Since orexin neurons in the lateral hypothalamus (LH) are involved in fear conditioning, and are inhibited by feeding, we determined whether they are involved in mood alterations observed in fear-conditioned fed GFAP-IR KO mice. We observed increased c-fos expression in orexin neurons of these animals, indicating that the lack of IR in astrocytes modifies the response of these neurons to food. Moreover, chemogenetic stimulation of LH astrocytes in fed GFAP-IR KO mice was sufficient to significantly attenuate their fear responses. These findings suggest that astrocytic insulin receptors modulate the responses of hypothalamic orexin neurons to food. We speculate that cellular mechanisms underlying eating disorders may involve insulin modulation of food valence through astrocytes.

## Introduction

Brain states such as hunger or fear interact with each other (Flavell et al., 2022) to modulate behavior, and are regulated by diverse signals, including hormones such as insulin, which is involved in regulation of feeding behavior (Volkow et al., 2011) acting as a satiety signal (VanderWeele, 1994), and modulation of food intake (Yang et al., 2015), exerting a central control of energy homeostasis (Scherer et al., 2021). In turn, bidirectional communication between neurons and astrocytes modulates different behavioral outputs involving subsets of neural networks (Oliveira and Araque, 2022). Astrocytes are ideally positioned to inform neuronal networks of changes in peripheral signals as their specialized cytoplasmic extensions known as end-feet connect the brain circulation with synaptic activity (Gourine et al 2011). In addition, astrocytes are sensitive to food (MacDonald et al., 2020), and act as insulin detectors (Garcia-Caceres et al 2016).

While the analysis of internal states with their pervasive influence on behavioral outputs allows a more comprehensive understanding of brain function (Flavell et al., 2022), many questions are still pending. For instance, it is generally assumed that neurons are the cellular base of internal states (Flavell et al., 2022), although glial cells such as astrocytes are also involved in brain states (Oliveira et 2022). Along this line, neurons are known to provide affective value to brain states (Allen et al., 2017), but astrocytes and other glial cells may also participate in the creation of motivation (Requie et al., 2022) and valence (Miyashita et al., 2023).

Building on the above observations, we determined whether this type of glial cell is involved in assigning a valence to food-related states, i.e. hunger and satiation, internal states driving feeding (Betley et al., 2015; Alhadeff et al., 2019). We used mice with tamoxifen-regulated ablation of the insulin receptor (IR) in astrocytes (GFAP-IR KO mice), a mouse that displays central and peripheral disturbances (González-García et al., 2021), including altered mood (Cai et al., 2018), and loss of astrocyte activation in response to insulin (Fernandez et al., 2022). Thus, we determined the influence of food in anxiety and fear, as proxies of the positive actions of food on mood. We observed that these mice showed negative affective responses to food that were normalized by chemogenetic activation of astrocytes in the lateral hypothalamus (LH), a region that is associated to food palatability (Castro et al., 2015), and is traditionally recognized as the feeding center of the brain.

Moreover, orexin neurons in the LH play a key role in modulating fear responses and overall mood (Sakurai et al., 1998). The activity of these neurons is regulated by both insulin (Silva et al., 2009) and food intake (González et al., 2016), and their interaction with hedonic hotspots suggests a role in assigning affective valence to food (Feillet et al., 2017). As already described, activation of orexin neurons was associated with fear responses. Notably, these fear responses were modulated by food availability, a process that depended on the functional state of LH astrocytes, as our DREADD-mediated modulation of astrocyte activity suggests.

Collectively, our findings indicate that astrocytes contribute to the assignment of emotional valence to food through insulin-mediated mechanisms, with LH astrocytes playing a specific role in this process by modulating the activity of orexin neurons.

## Results

### Astrocytes are involved in insulin modulation of food valence

To measure the positive valence of food in mice we determined its lowering effects on anxiety levels as a proxy of improved mood, using the elevated plus maze (EPM) and the open field tests (OFT). We first conditioned wild type mice to palatable food and drink (Figure 1A,B and Suppl Fig 1A,B), and then assessed whether their anxiety was reduced using the elevated plus maze (EPM) and open field tests (Figure 1C,D). Then we tested conditioned GFAP-IR KO mice. While both control and GFAP-IR KO mice were conditioned by palatable food, as determined by more time spent in the food/drink chamber (Figure 1E), the rewarding properties of palatable food were observed only in control littermates. Reduced anxiety, as determined by increased time spent in the open arms of the elevated plus maze (EPM, Figure 1F) and in the center of the open field (Figure 1G), was seen only in control mice, but not in GFAP-IR KO mice (Figure 1F,G). Thus, while the conditioning properties of food are intact in mice lacking IR in astrocytes, their rewarding properties are lacking.

**Figure 1.**
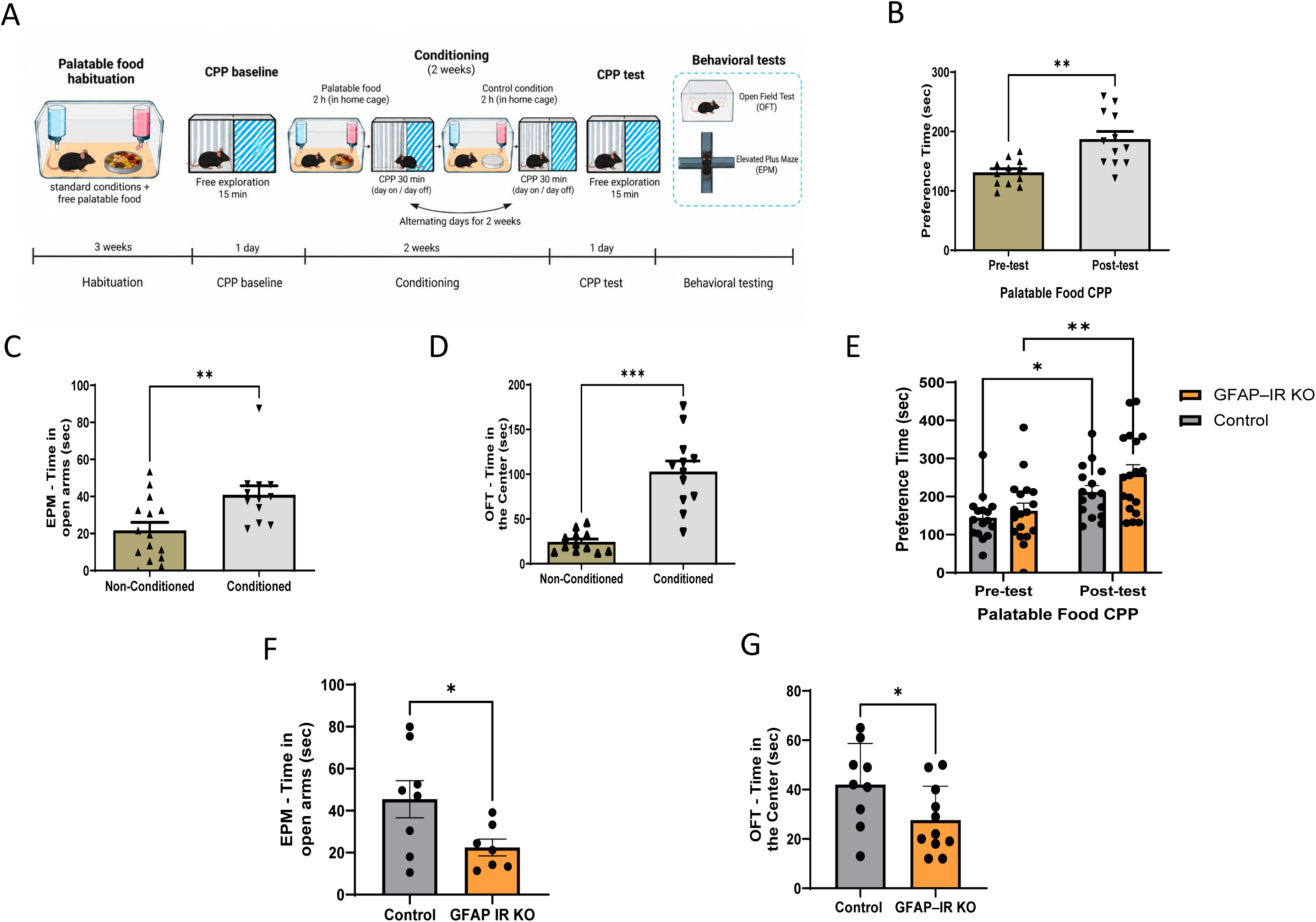
Mice conditioned with a palatable food show improved mood. **A**, Food-conditioning protocol was followed by behavioral testing of conditioned mice. CPP: conditioned place preference test. **B,** Place preference was observed in conditioned wild-type mice before (pre-test) and after (post-test) food conditioning (n = 12; paired two-tailed Student’s t-test, p = 0.0012). **C,** Food-conditioned wild-type mice spent more time in the open arms of the elevated plus maze (EPM) than unconditioned mice (unconditioned, n = 15; conditioned, n = 12; two-tailed Mann–Whitney U test, p = 0.0077). **D,** After conditioning, wild-type mice spent more time in the center of the open field arena (n = 12; paired two-tailed Student’s t-test, p = 0.0001). **E,** Place preference was observed in GFAP-IR KO mice (n = 18) and control littermates (n = 16) after conditioning with palatable food (mixed-effects model [REML] followed by Šidák’s multiple comparisons test). **F,** Conditioned GFAP-IR KO mice (n = 7) spent less time in the open arms of the EPM compared with control littermates receiving oil (n = 8; two-tailed Welch’s t-test, p = 0.0394). **G,** Conditioned GFAP-IR KO mice (n = 11) spent less time in the center of the open field arena compared with control littermates (n = 9; unpaired two-tailed Student’s t-test, p = 0.0489). Data are presented as mean ± SEM. *p < 0.05, **p < 0.01, ***p < 0.001.

Furthermore, feeding GFAP-IR KO mice with standard diet appears to deteriorate their mood. Thus, using fear responses as a proxy of changes in mood, we found that when *ad lib* fed GFAP-IR KO mice were conditioned using classical fear-conditioning protocols (Figure 2A), they showed increased fear responses compared to vehicle-treated siblings (Figure 2B). Even during the training phase, fed GFAP-IR KO showed significantly higher freezing (Figure 2B, Cs-) suggesting that these mice display altered fear behavior during conditioning. This agrees with previous observations of disrupted mood homeostasis in GFAP-IR KO mice (Cai et al., 2018; Zegarra-Valdivia et al., 2022). To rule out possible unspecific actions of tamoxifen on behavior, we repeated these experiments in IR^f/f^ mice injected with a GFAP-Cre virus in the tail (BIR GFAP Cre mice, Figure 2C), that present brain-wide reduction of IR in GFAP astrocytes (Suppl Fig 1C,D). These mice, when fed, showed the same enhanced fear responses as fed GFAP-IR KO mice (Figure 2D).

**Figure 2.**
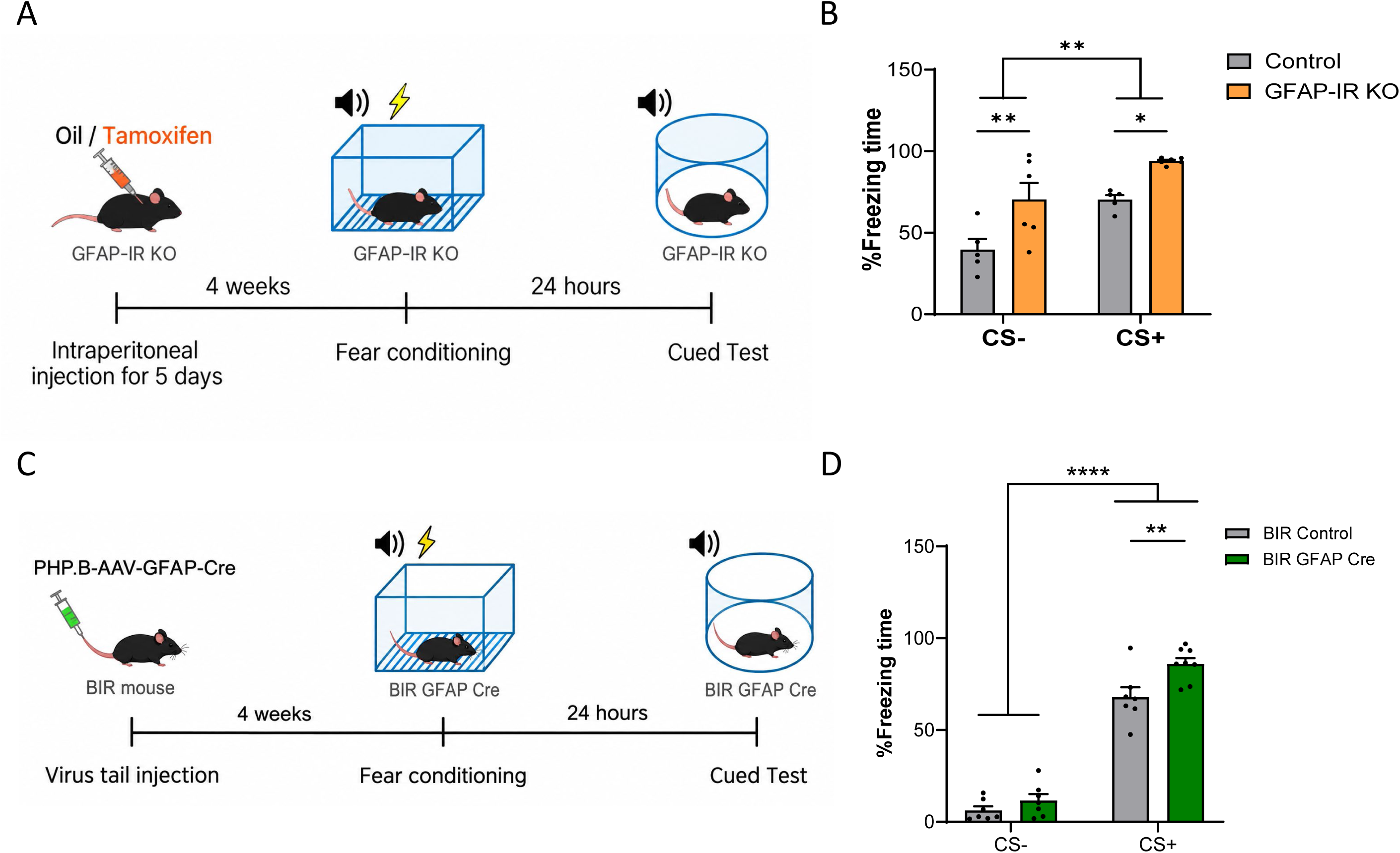
The insulin receptor (IR) in astrocytes modulates fear learning. **A**, Time-course of experimental procedures followed in fear conditioning of GFAP-IR KO mice. **B,** GFAP-IR KO mice presented a higher percentage of freezing time during CS− and CS+ periods in the cued fear test compared with control littermates (Control, n = 5; GFAP-IR KO, n = 6; two-way repeated-measures ANOVA followed by Šidák’s multiple comparisons test; CS−, p = 0.0068; CS+, p = 0.0366). **C,** Time-course of experimental procedures in mice with viral-mediated knockdown of IR in astrocytes. **D,** BIR GFAP Cre mice presented a higher percentage of freezing time during CS+ presentations compared with BIR Control mice (BIR Control, n = 7; BIR GFAP Cre, n = 8; mixed-effects model [REML] followed by Šidák’s multiple comparisons test; CS+, p = 0.0035). Data are presented as mean ± SEM, with individual points representing mice. *p < 0.05, **p < 0.01, ****p < 0.0001.

We then examined the role of the lateral hypothalamus (LH)-Locus coeruleus (LC) circuit involved in freezing behavior (Soya et al., 2017), in fear-conditioned GFAP-IR KO mice, by determining c-fos immunostaining, which is a commonly used indicator of neuronal activation (Sundquist and Nisenbaum, 2005). Both LH orexin neurons and LC tyrosine hydroxylase (TH) neurons, heavily innervated by orexin neurons (Date et al., 1999), showed increased c-fos staining in fear-conditioned GFAP-IR KO mice (Figure 3A-C). This pattern of increased activation was also observed in orexin and LC TH neurons of fear-conditioned BIR GFAP Cre mice (Suppl Fig 2A-C). To further confirm the role of orexin neurons of GFAP-IR KO in this exaggerated fear behavior, we used optogenetic activation of these neurons using orexin-ChR2 viruses in fear-conditioned GFAP-IR KO mice. Upon light stimulation, greater fear responses were elicited in GFAP-IR KO mice than in littermates (Figure 4), corroborating the involvement of these neurons in the observed behavior.

**Figure 3.**
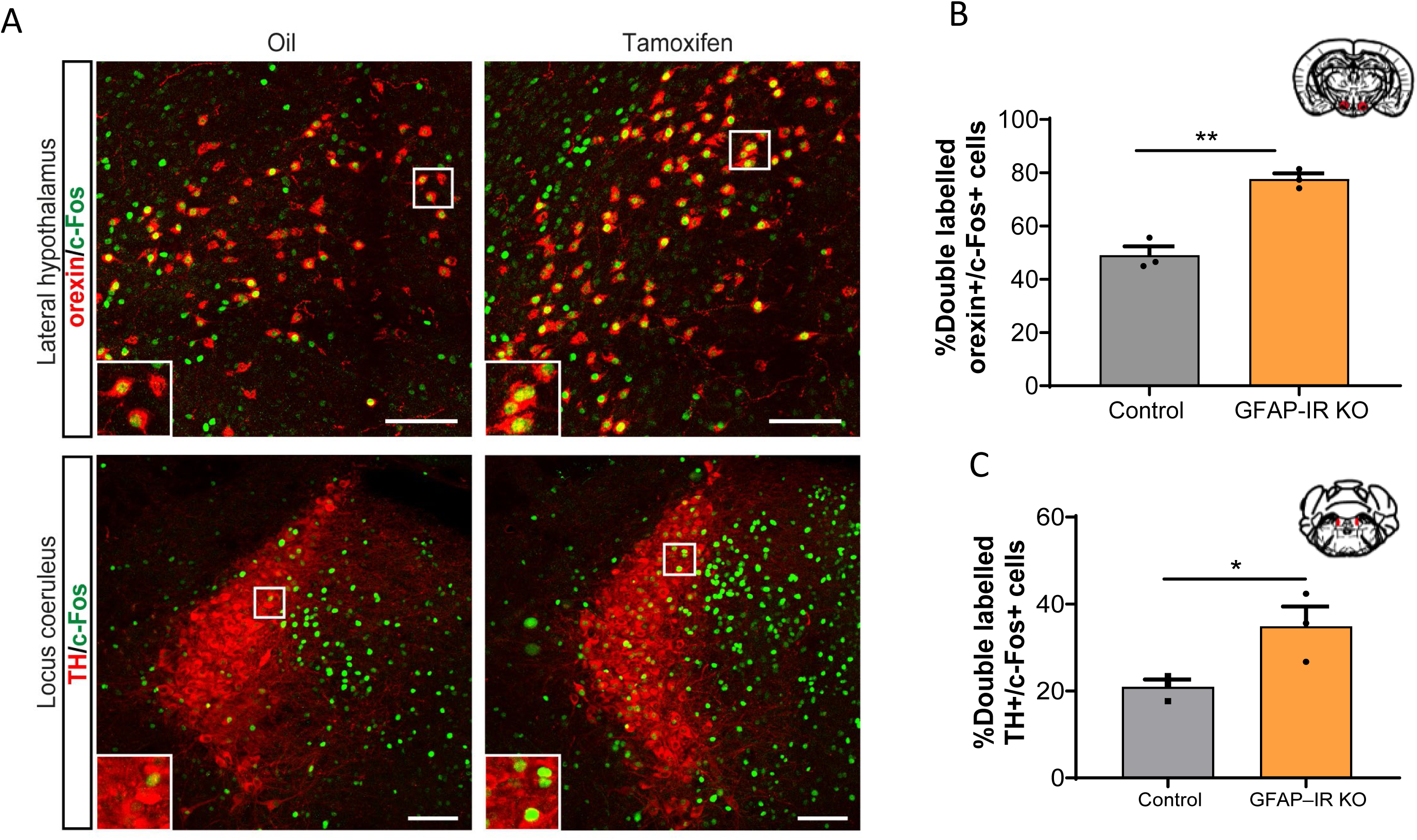
Astrocytic insulin receptor signaling modulates c-Fos expression in fear-related nuclei following fear conditioning. **A**, Representative micrographs showing double immunostaining for c-Fos (green) and orexin (red) in the lateral hypothalamus (upper panels), and c-Fos (green) and tyrosine hydroxylase (TH; red) in the locus coeruleus (LC; lower panels) of oil-treated control mice (left) and tamoxifen-treated GFAP-IR KO mice (right) following fear conditioning. **B,** The percentage of orexin-positive neurons coexpressing c-Fos was significantly increased in GFAP-IR KO mice compared with controls (Control, n = 3; GFAP-IR KO, n = 3; unpaired two-tailed Student’s t-test, t(4) = 7.308, p = 0.0019). **C,** The percentage of TH-positive neurons coexpressing c-Fos in the LC was significantly increased in GFAP-IR KO mice compared with controls (Control, n = 3; GFAP-IR KO, n = 3; unpaired two-tailed Student’s t-test, t(4) = 2.876, p = 0.0452). Data are presented as mean ± SEM, with individual points representing mice. Scale bars: 100 µm. *p < 0.05, **p < 0.01.

**Figure 4.**
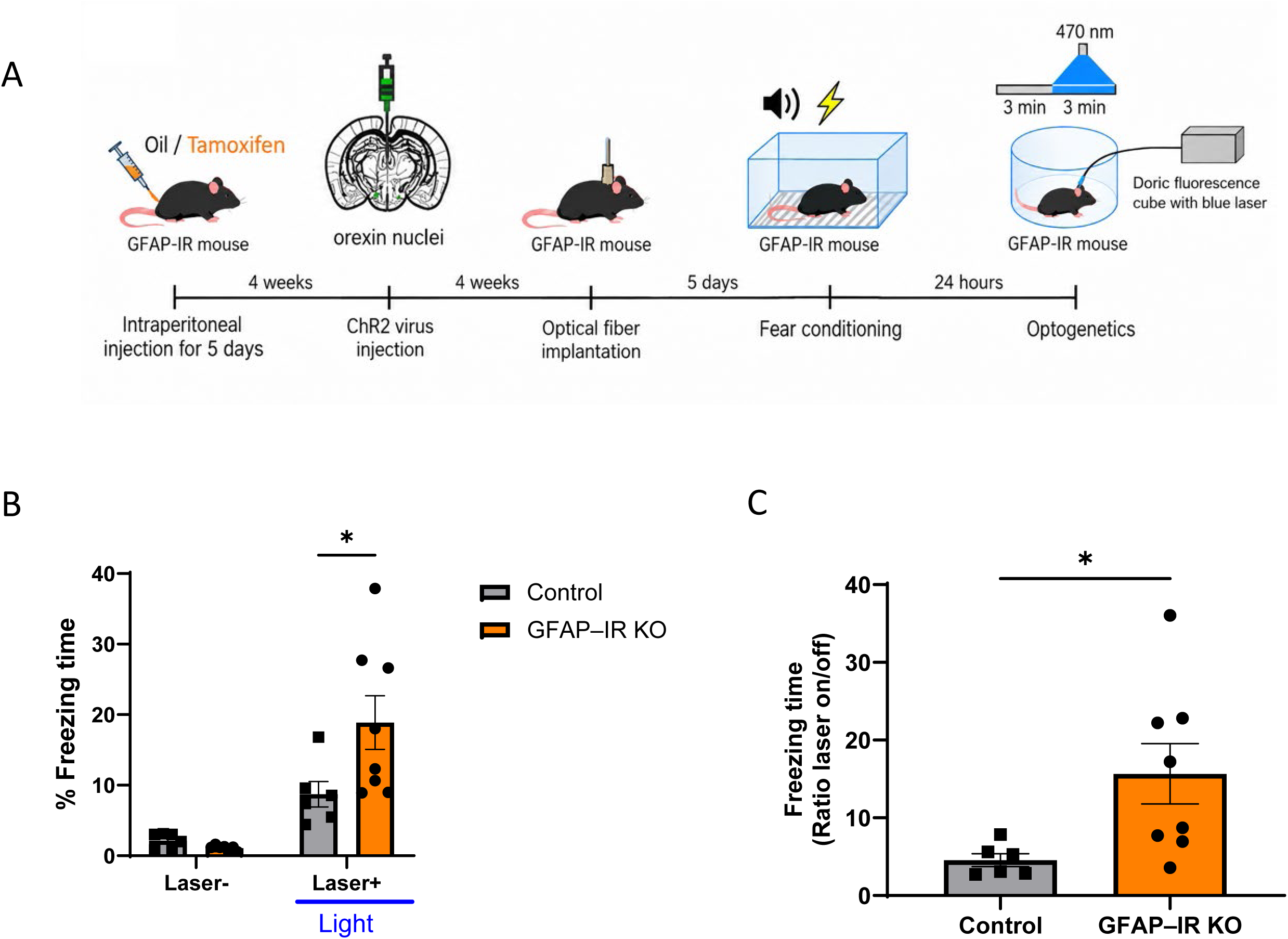
Optogenetic activation of orexin neurons in GFAP-IR KO mice increases freezing in the cued fear test. **A**, Time-course of experimental procedures followed in optogenetic experiments. **B,** Optogenetic activation of orexin neurons in tamoxifen-injected GFAP-IR KO mice enhances freezing responses in the cued test during laser stimulation (Control, n = 6; GFAP-IR KO, n = 8; repeated-measures mixed-effects model [REML] followed by Šidák’s multiple comparisons test; laser ON, p = 0.0142). **C,** Freezing time was also higher in the GFAP-IR KO group when expressed as the laser ON/OFF ratio (Control, n = 6; GFAP-IR KO, n = 8; two-tailed Welch’s t-test, t(7.637) = 2.792, p = 0.0246). Data are presented as mean ± SEM, with individual points representing mice. *p < 0.05.

To assess whether food carries a negative valence in GFAP-IR KO mice, we examined fear responses in fasted animals, based on prior evidence suggesting an interaction between hunger and fear (Burnett et al., 2016). We observed that under fasting, GFAP-IR KO mice show reduced fear as compared to littermates (Figure 5A). Conversely, food reduced fear expression both in littermates (Figure 5A) and wild type controls (Suppl Fig 1E), as expected. To confirm these observations, we determined anxiety levels in fasted GFAP-IR KO mice and, again, observed reduced anxiety in the EPM, as compared to littermates (Figure 5B).

**Figure 5.**
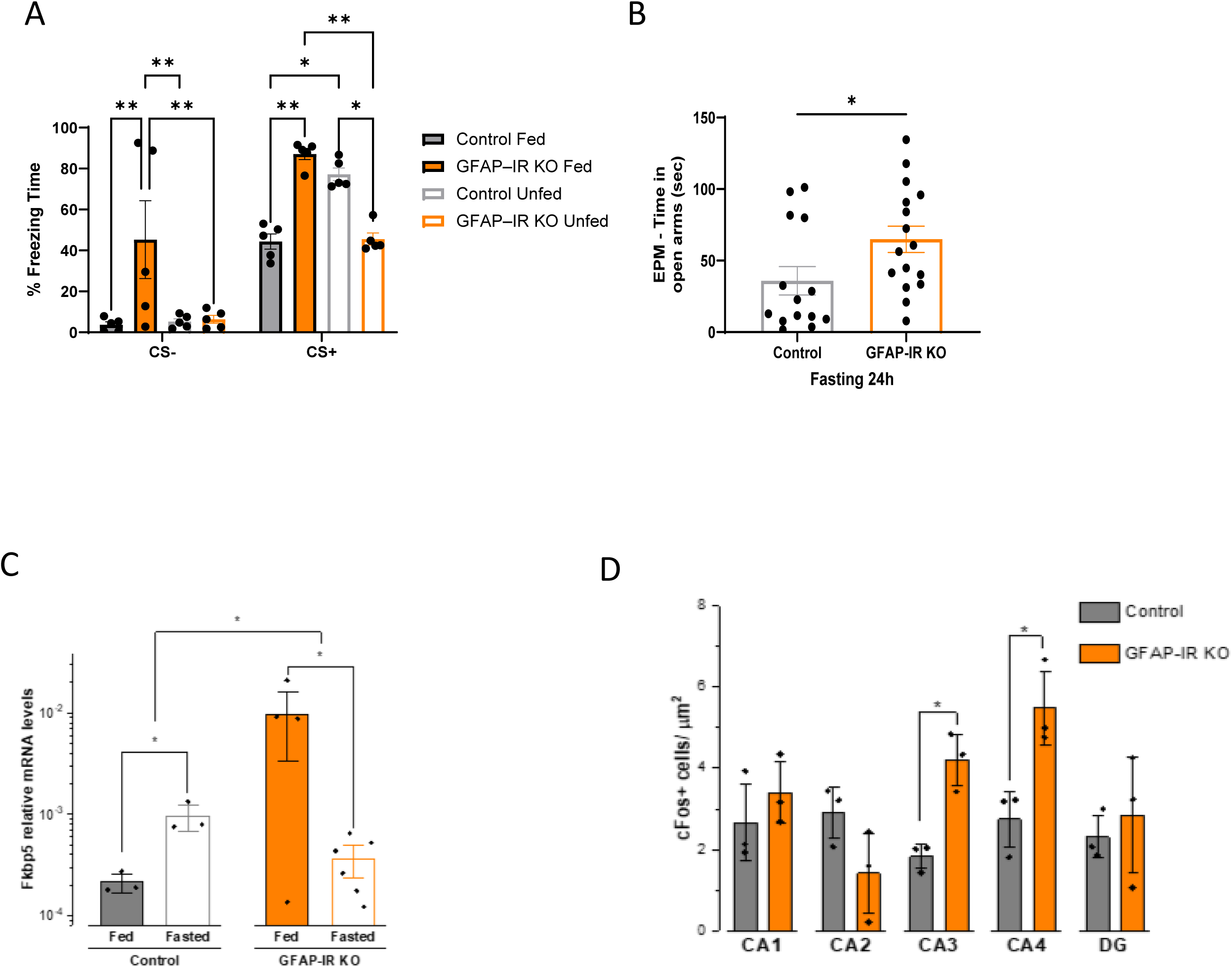
Food shows a negative valence in GFAP-IR KO mice. **A**, Opposite effects of fasting on conditioned freezing in control and GFAP-IR KO mice. Freezing was reduced in fasted GFAP-IR KO mice, whereas it increased in fasted control littermates (Control Fed, n = 5; Control Fasted, n = 5; GFAP-IR KO Fed, n = 5; GFAP-IR KO Fasted, n = 5). Two-way repeated-measures ANOVA followed by Šidák’s multiple comparisons test. A complementary three-way repeated-measures ANOVA revealed a significant genotype × nutritional state interaction (F(1,16) = 32.99, p < 0.0001). **B,** Following 24 h of fasting, GFAP-IR KO mice spent more time in the open arms of the elevated plus maze (EPM) than control littermates (Control, n = 14; GFAP-IR KO, n = 16; two-tailed Mann–Whitney U test, U = 57, p = 0.0218). **C,** Hippocampal FKBP5 expression was differentially regulated by nutritional state in control and GFAP-IR KO mice (n = 3–6 mice per group; ANOVA followed by Tukey’s multiple comparisons test; *p < 0.05). **D,** Quantification of c-Fos-positive cells per µm² in different hippocampal regions following fear conditioning in fed and fasted control and GFAP-IR KO mice. Fed GFAP-IR KO mice exhibited greater c-Fos-positive cell density in CA3 and CA4 than fasted GFAP-IR KO mice (two-tailed Student’s t-test; *p < 0.05). CA, cornu ammonis; DG, dentate gyrus. Data are presented as mean ± SEM, with individual points representing mice. *p < 0.05, **p < 0.01, ***p < 0.001.

### Hippocampal FKBP5 and oxidative stress participate in insulin actions through astrocytes

We next examined possible mechanisms underlying the effect of impaired IR activity in astrocytes on food valence. Responses to stress are modulated by hippocampal levels of the food-sensitive FK506 binding protein 51 (FKBP5) (Scharf et al., 2011), a co-chaperone of the glucocorticoid receptor (Criado-Marrero et al., 2019), and a biomarker of its activity (Binder, 2009). Levels of FKBP5 are also related to insulin resistance (Strączkowski et al., 2022), which is present in the brain of GFAP-IR KO mice (Garcia-Caceres et al., 2016). Thus, we measured its mRNA in the hippocampus of fed and fasted GFAP-IR KO mice and their corresponding littermates. As shown in Figure 5C, modulation by food of the expression of hippocampal FKBP5 was altered in GFAP-IR KO; increased in fasted controls (as reported), and decreased in fasted GFAP-IR KOs, paralleling fear responses (see Figure 5A, right histograms). That fed GFAP-IR KO mice showed increased expression of hippocampal FKBP5 suggests that the activity of the hippocampal glucocorticoid receptor is increased (Binder, 2009), supporting the observation of heightened levels of fear stress in fear-conditioned fed GFAP-IR KO mice. Indeed, hippocampal neuronal activation in fear-conditioned mice showed that in fed GFAP-IR KO mice the CA3 and CA4 regions were significantly more activated, as determined by c-fos staining, compared to unfed GFAP-IR KO mice (Figure 5D). No differences in hippocampal c-fos staining were seen in unfed vs fed GFAP-IR KO or control mice, under basal conditions (Suppl Fig 3A).

Furthermore, since food modulates brain oxidative stress (Hazzaa et al., 2020) and GFAP-IR KO mice show increased brain oxidative stress (Fernandez et al., 2022), we determined whether behavioral responses to food would be modulated by administration of the anti-oxidant N-Acetyl-cysteine (NAC, 1% in the drinking water, 1 month), since NAC has previously been shown to modulate brain activity in GFAP-IR KO mice (Fernandez et al., 2022). NAC affected the responses to fear conditioning of both littermates and GFAP-IR KO mice in a differential way (Figure 6A), resulting in abolition of differences between the two groups. Significantly, NAC eliminated food-dependent fear responses in both groups.

**Figure 6.**
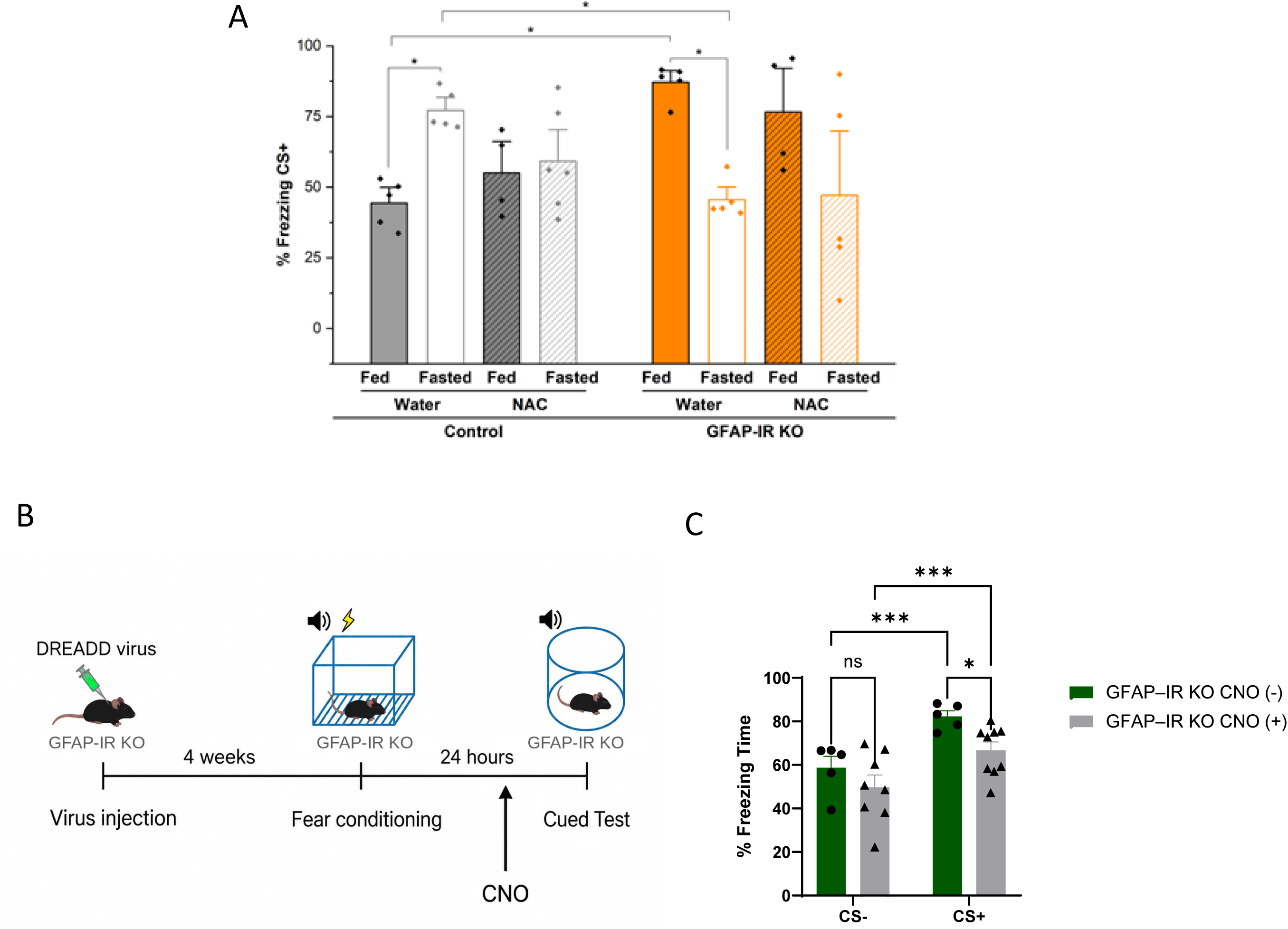
Mechanisms underlying negative valence of food in GFAP-IR KO mice. **A**, Treatment with N-acetylcysteine (NAC, 1% in drinking water for one month) attenuated freezing behavior in fasted control mice and fed GFAP-IR KO mice, reducing the behavioral differences associated with genotype and nutritional state (n = 5–6 mice per group; ANOVA followed by Tukey’s multiple comparisons test; *p < 0.05), *p < 0.05). **B,** Schematic representation of the experimental timeline for chemogenetic modulation of astrocytes in GFAP-IR KO and wild-type mice, including DREADD expression, CNO administration, and subsequent fear-conditioning assessment. **C,** Chemogenetic activation of lateral hypothalamic astrocytes in fed GFAP-IR KO mice reduced freezing responses during CS+ presentations following CNO administration (CNO−, n = 5; CNO+, n = 9). Mixed-effects analysis (REML) revealed a significant effect of stimulus period (F(1,11) = 59.00, p < 0.0001), whereas neither the main effect of CNO treatment (F(1,12) = 3.976, p = 0.0694) nor the stimulus period × treatment interaction (F(1,11) = 1.219, p = 0.2930) was significant. Fisher’s uncorrected LSD comparisons indicated reduced CS+ freezing in CNO-treated mice (p = 0.0322). Data are presented as mean ± SEM, with individual points representing mice. *p < 0.05; ***p < 0.001.

### LH astrocytes are involved in assigning food valence

Insulin reportedly modulates orexin neurons in the lateral hypothalamus (LH) (Silva et al., 2009), which project to the ventral tegmental area (VTA) and nucleus accumbens (NAc). Since these two regions are implicated in reward processing and the hedonic valuation of food (Fadel and Deutch, 2002) (Sakurai and Mieda, 2011) (Harris et al., 2005), and therefore are influenced by food intake (González et al., 2016), we investigated the potential role of astrocyte–orexin neuron cross-talk in assigning affective value to food. We stimulated LH astrocytes in fed GFAP-IR KO mice using hM3Dq DREADD viruses as described (Fernández de Sevilla et al., 2022), and determined fear responses after administration of CNO. Chemogenetic stimulation of LH astrocytes significantly attenuated fear responses in fed GFAP-IR KO mice, suggesting that astrocyte-orexin neurons cross-talk participates in exaggerated fear responses in these mice (Figure 6B,C). However, inhibition of LH astrocytes in fed wild type mice with hM4Di DREADD viruses did not alter fear responses (Suppl Fig 3D), suggesting that altered activity of LH astrocytes *per se* is not enough to disturb fear responses in fed mice.

## Discussion

These results indicate that astrocytes participate in modulation of food valence in a process affecting different brain sites and mechanisms, including modulation of orexin neurons in the lateral hypothalamus, and of brain levels of oxidative stress. This suggests that astrocytes constitute a cellular basis of valence in mammals, suggesting that glial cells maintain this role throughout evolution as ensheathing glia in flies are involved in valence transmission (Miyashita et al., 2023). Moreover, the observation that GFAP-IR KO mice develop conditioned associations with palatable food but do not exhibit mood improvement, suggests that the rewarding properties of food — independent of hunger-driven conditioning, are modulated by astrocytic insulin receptors (IR). This finding highlights a functional dissociation between reward and metabolic signaling pathways at the level of brain circuitry (Nieh et al., 2015). These results may also suggest that maladaptive processing of stimulus valence underlying numerous psychiatric and metabolic conditions (Chen, 2023), such as *anorexia nervosa* (Kaye, 2008) or obesity (Mazzone et al., 2020) may include insulin-mediated astrocyte influence on neuronal networks (Kwak et al., 2020; Oliveira and Araque, 2022; Noh et al., 2023).

In response to food, insulin is released by the pancreas (Rutter, 2001), and modulates feeding behavior, acting as a satiety signal (Berthoud and Baettig, 1974). Since GFAP IR KO mice show lower sensitivity to circulating insulin (Fernandez et al., 2022), the positive feedback information produced by food consumption (Berthoud et al., 2011) seems to be impaired in them, as they show reduced satiety (Garcia-Caceres et al., 2016). This deficit may also interfere with the rewarding properties of food since the hedonic component of palatable food is dissociated from its conditioning effects in GFAP IR KO mice, which developed preference for palatable food in the absence of noticeable improvement in their mood. This disconnection may relate to the depressive-like behavior shown by GFAP-IR KO mice (Cai et al., 2018; Zegarra-Valdivia et al., 2022), as depression is associated to altered function of the reward circuit (Russo and Nestler, 2013). Further, GFAP-IR KO mice show a wide panoply of brain disturbances, including altered brain glucose-sensing (Garcia-Caceres et al., 2016), neurovascular uncoupling (Fernandez et al., 2022), and reduced brain glucose uptake (Fernandez et al., 2017). Collectively, these disturbances may contribute to the observed food-dependent phenotype of these animals.

Parallel changes in hippocampal FKBP5 expression and fear responses in fed and unfed states both in controls and GFAP-IR KO mice indicates that this glucocorticoid receptor chaperone that intervenes in responses to stress (Criado-Marrero et al., 2019) and is increased by fasting (Scharf et al., 2011), is acting as a biomarker of glucocorticoid activity (Binder, 2009), faithfully reflecting stress levels. Since fasting GFAP-IR KO mice did not show increased hippocampal FKBP5, as seen in controls, we speculate that loss of astrocytic insulin sensitivity disturbs neuronal responses to stress at the hypothalamic paraventricular nucleus, that orchestrates stress responses to fasting (Melnick et al., 2020). Altered coupling between nutritional state and stress-related signaling was further reflected by elevated hippocampal FKBP5 expression in fed GFAP-IR KO mice.

Mice lacking IR in astrocytes showed increased brain ROS that is amenable to treatment with the anti-oxidant NAC (Fernandez et al., 2022). Since NAC abolished not only the differences in fear responses between GFAP-IR KO mice and their littermates, but also the effects of food on fear responses regardless of the presence or absence of IR in astrocytes, we consider that ROS by its own is important in assigning food valence. Thus, through interference with ROS levels in the brain (Fernandez et al., 2022), lack of IR in astrocytes interferes with food valence too. This would be an additional mediator of the actions of astrocytic IR on food valence.

Another potential downstream target of astrocytes in regulation of food valence is LH neurons. The role of the LH in modulating the rewarding properties of food was described long ago (Delgado and Anand, 1952). Our data would support the inclusion of LH astrocytes in the modulation of food valence by directly or indirectly affecting orexin neurons since these neurons are more activated in response to fear-learning in GFAP-IR KO mice and stimulation of LH astrocytes with DREADD in fed GFAP-IR KO mice reduced fear responses to food. Conceivably, as previously seen in enhanced orexinergic activity in mice with exaggerated fear responses (Fernandez de Sevilla et al., 2022), the mechanism whereby orexin neurons are overactivated in fed GFAP-IR KO mice might be through lower inhibitory input on them. Indeed, insulin stimulates GABA activity (Hammoud et al., 2021), and in LH GABA neurons this effect could be indirect through astrocytes, as insulin modulates gliotransmitters (Cai et al., 2018), that can modulate neuronal activity (Araque et al., 2014).

Alternatively, since in GFAP-IR KO mice insulin no longer stimulates ATP release (Cai et al., 2018), and Ca^++^ transients (Fernandez et al., 2022), it may also directly impair GABA release by astrocytes (Kwak et al., 2020). Thus, either indirectly through LH GABA neurons or through reduced GABA release by astrocytes, the result would be reduced inhibition on orexin neurons (Figure 7). In turn, low inhibitory shunting tone on orexin neurons due to the lack of astrocytic insulin signaling would overexcite them during fear-conditioning (Figure 7), with downstream overactivation of LC neurons, which are important mediators of freezing responses (Soya et al., 2017). The fact that insulin has been shown to directly modulate orexinergic activity (Silva et al., 2009) may compound the role of this hormone on orexin signaling. Other peptide signals have been shown to directly stimulate orexin and indirectly, through GABA neurons, inhibit it (Furutani et al., 2010), so this mechanism is plausible. Potential molecular mediators include the nutrient sensor AMPK (González et al., 2020). Hypothalamic AMPK regulates food intake and insulin regulates it acting at multiple hypothalamic sites (Minokoshi et al., 2004). Conceivably, AMPK in hypothalamic astrocytes in the LH may be a target of insulin to regulate food behavior.

**Figure 7.**
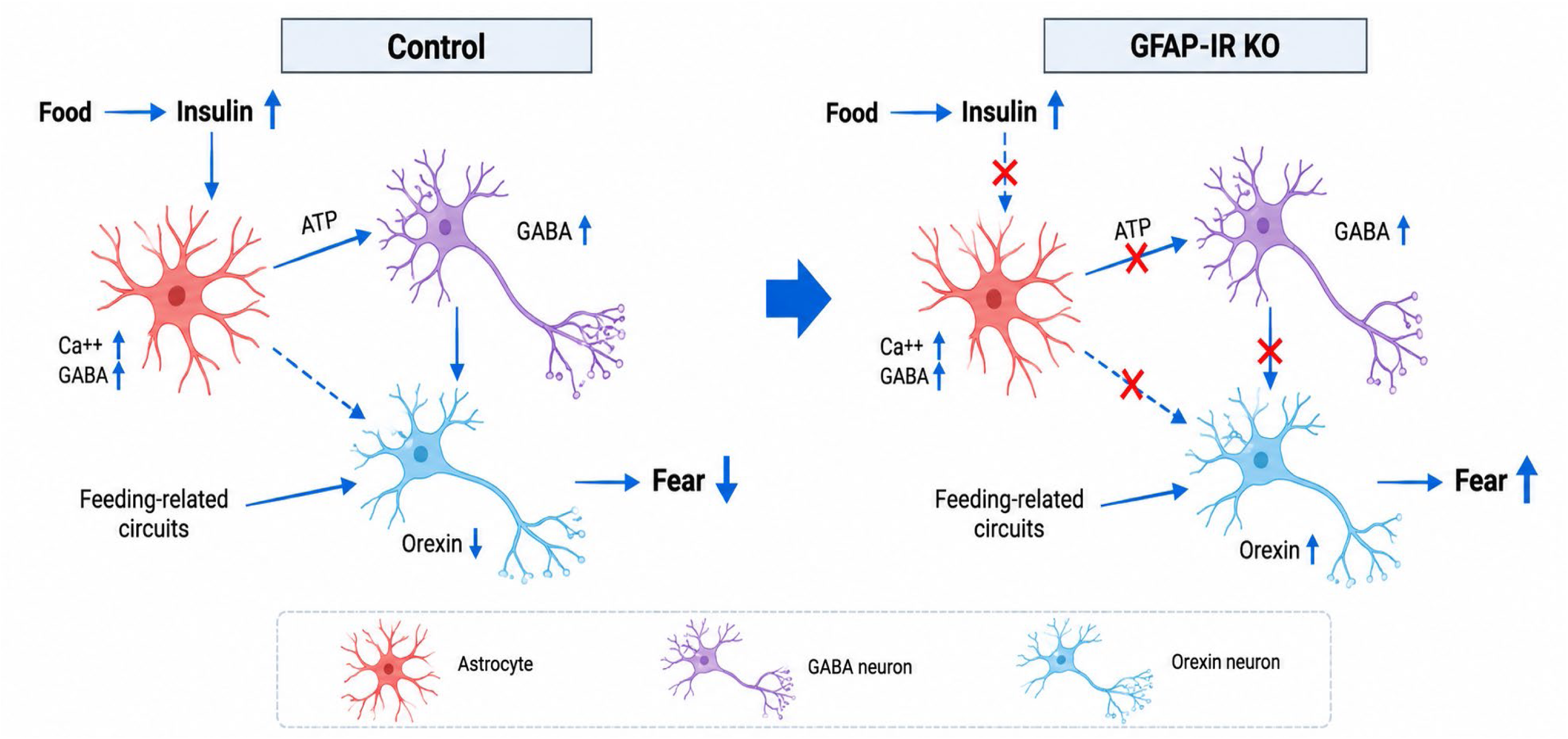
Food inhibits orexin activity in part through astrocytes in an insulin-dependent manner. In response to food intake, insulin is released by the pancreas and reaches the brain, where it contributes to the regulation of feeding behavior. We propose that astrocytes in the lateral hypothalamus (LH), a key region involved in feeding regulation, respond to insulin and promote GABA release either directly or indirectly through local GABAergic neurons. GABA, in turn, tonically inhibits orexin neurons through GABA_A receptors (Toossi et al., 2016). Orexin neuronal activity is inhibited by food intake (González et al., 2016), potentially involving additional feeding-related circuits (Broberger et al., 1998; Hsu et al., 2015) as part of the homeostatic response to food (Inutsuka et al., 2014). In GFAP-IR KO mice, impaired astrocytic insulin signaling may reduce GABAergic inhibition of orexin neurons, resulting in increased orexin activity and enhanced conditioned fear responses.

A limitation of this study is that we have not ruled out the possibility that hunger itself may differentially modulate sensory processing (Tallon-Baudry, 2023) in GFAP-IR KO mice. Hence, further studies should determine whether hearing and/or pain perception involved in fear-learning are distinctly altered in fasted GFAP-IR KO mice. An additional limitation is that we used the level of expression of negative affect as an indirect measure of positive valence of food. Evaluation of positive emotions in mice, although challenging, would constitute a more direct assessment. Also, other brain areas such as prefrontal cortex and amygdala, where astrocytes also play a role in assigning valence, should be included as they are involved in valence assignment and fear responses, respectively.

In summary, our observations widen the actions of insulin in the brain and assign a role of hypothalamic astrocytes, interacting with hypothalamic neurons, in modulating valence to food. Also, levels of oxidative stress in brain appear to modulate food valence.

## Materials and Methods

### Animals

Mice were used according to Animal Research Reporting of in vivo Experiments (ARRIVE), and this study is reported in line with the guidelines (Zegarra-Valdivia et al., 2022). Transgenic mice with tamoxifen-regulated deletion of IR in astrocytes (GFAP-IR KO mice) were obtained as described (Garcia-Caceres et al., 2016; Noriega-Prieto et al., 2021) crossing IR ^f/f^ (B6.129S4(FVB)-Insr ^tm1Khn^/J RRID:IMSR, Jackson labs; stock number 006955) with hGFAP-CreER ^T2^ mice (C57B&/6xSJL/J mix background Jackson Labs, stock number: 012849). To knock down the target gene, tamoxifen was administered to 2-months old mice for 5 days (75 mg/kg, Sigma, intraperitoneally), as described (Hirrlinger *et al*., 2006), and animals were used one month later. Controls were littermates receiving the vehicle (corn oil). GFAP-IR KO mice display reduced IR mRNA levels in brain, as reported by Garcia-Caceres *et al*. (2016). GFAP-IR KO mice show brain IGF-IR levels similar to wild type mice. Mice were housed in standard cages (48 × 26 cm ^2^) with 5 mice per cage and kept on a light-dark cycle (12-12 h, lights on at 8 am) at constant temperature (22°C) and humidity, and with food (pellet rodent diet) and water *ad libitum.* All experimental protocols were performed during the light cycle and followed European guidelines (86/609/EEC & 2003/65/EC, European Council Directives). Studies were approved by the respective local Bioethics Committees (Government of the Community of Madrid, PROEX 193.4/20 (2020) and UPV M20_2021_168 (2021). Animals were not randomized and were used in a sex-balanced manner throughout. Potential confounders were not accounted for. Each experimenter took account of group allocation under study. All efforts were made to reduce harm and the number of animals. Mice were handled for three days prior to any experimental manipulations and familiarized with behavioral arenas to minimize novelty stress. Mice were deeply anesthetized with pentobarbital prior to sacrifice, when needed. Sample sizes were kept as small as possible to comply with current animal reduction policies. No adverse events were expected, nor found. End-point measures included checking reflexes in deeply anesthetized animals prior to culling.

### Reagents

The following reagents were used: N-acetyl-cysteine (NAC; Sigma, Darmstadt, Germany), and Tamoxifen (Sigma). NAC was administered at a concentration of 1% in the drinking water for one month. Antibodies used were: ß-actin (Sigma cat. no. A5316), GFAP (Dako, Glostrup, DK, cat. no. Z0334), Insulin receptor ß (Santa Cruz cat. no. 23413), NeuN (Proteintech, cat. no. 26975-1-AP). Antibodies against c-fos, TH, and orexin, and Hoechst 33342 (DAPI) were purchased from Invitrogen. FKBP5 inhibitor was given ip at 20mg/kg and behaviour was tested 2 hours later.

### Viral constructs

For optogenetic experiments we used ChR2 (AAV8-HCRT-CHR2-EGFP; 1.33 x 10^13^ viral infective units/ml) viruses injected into the orexin nuclei of GFAP-IR KO mice or littermates, as explained in detail before (Fernández de Sevilla et al., 2022). To modulate the activity of GFAP astrocytes in the lateral hypothalamus of GFAP-IR KO mice and in wild type mice we used inhibitory and stimulatory DREADD viruses, as explained before. Clozapine N-Oxide (CNO, 2mg/kg dissolved in saline 0.9%) was administered ip 24 h after fear-conditioning, and 40 min later behavior was assessed. For inactivation of IR in GFAP^+^ astrocytes of adult mice, we injected in the tail of IR^f/f^ mice (3 months-old), a PHP B-AAV-GFAP-Cre-EGFP (1.33 x 10^13^ viral infective units/ml) virus (AAV-GFAP). In controls we injected in the tail a PHP B-AAV-CMV-EGFP (1.3 x 10^13^ viral infective units/ml) virus (AAV-CMV-Control). One month later we submitted them to fear-learning (see below).

For astrocyte-directed chemogenetic activation, an excitatory hM3Dq construct (ssAAV-8/2-hGFAP-hM3D(Gq)-mCherry-WPRE-hGHp(A); laboratory inventory v97-8) was used in GFAP-IR KO mice. For astrocyte-directed inhibition, an inhibitory hM4Di construct (ssAAV-8/2-hGFAP-hM4D(Gi)-mCherry-WPRE-hGHp(A); inventory v103-8) was used in wild-type mice. The chemogenetic procedures were adapted from prior work (Fernández de Sevilla et al., 2022; Zegarra-Valdivia et al., 2026), with astrocyte-directed DREADD vectors.

### Surgery

Mice were anesthetized with isoflurane (Zoetis) administered through a nose mask (David Kopf Instruments, France) and positioned in a stereotaxic frame (Stoelting Co.) on a heating pad. Their eyes were protected from light during surgery. Viral vectors were bilaterally injected into the lateral hypothalamus (LH), targeting the orexin-containing region, using a 5-µL Hamilton syringe and the following stereotaxic coordinates relative to bregma: anteroposterior (AP), −1.4 mm; mediolateral (ML), ±0.9 mm; and dorsoventral (DV), −5.4 mm.

For DREADD-mediated modulation of LH astrocytes, 300 nL of viral suspension was infused per injection site at a rate of 100 nL/min. The Hamilton syringe was left in place for an additional 10 min after infusion to minimize viral reflux. Following stereotaxic surgery, mice were allowed a 4-week recovery and viral expression period before undergoing behavioral testing.

### Behavioral tests

#### Conditioned place preference induced by palatable food and drink

Conditioned place preference (CPP) was used to assess the motivational effects of palatable food and drink, following the same protocol reported previously (Zegarra-Valdivia et al., 2026), but with modifications. Briefly, before CPP testing, mice were familiarized with the palatable foods and drink used as rewarding stimuli, including sunflower seeds, biscuits, and sweetened condensed milk diluted in water, as a habituation phase. After 3 weeks, a pre-test was done. The apparatus comprised two conditioning compartments with distinct visual cues (a white chamber with black dots and a white chamber with gray vertical stripes) separated by a neutral black central compartment. At pretest, mice spent 5 min in each compartment for habituation and then explored all compartments freely for 15 min. Baseline time spent in each compartment was used to assign the less-preferred compartment to the reward-paired condition and the preferred compartment to the unpaired condition. Conditioning was conducted over 14 days with alternating paired and unpaired sessions. In paired sessions, animals had 2 h access to palatable food or drink in their home cages, followed by 30 min confinement to the paired compartment. Palatable stimuli included sunflower seeds, biscuits, and sweetened condensed milk diluted in water. In unpaired sessions, the same containers were present in the home cages but empty, followed by 30 min confinement to the alternative chamber of CPP. Standard chow and water remained available ad libitum, without food restriction. Preliminary experiments examined food and drink conditioning separately; following comparable conditioning responses, both were used in the main experimental paradigm. On day 15, mice were allowed 15 min of free exploration without presentation of the palatable stimulus; conditioned preference was assessed as the change in time spent in the reward-paired compartment from pretest to post-test. Both sexes were included. GFAP-IR KO mice were tested after one month of tamoxifen induction. Subsequent open-field and elevated-plus-maze testing followed the sequence indicated in Fig. 1, with 2 h exposure to palatable stimuli before each behavioral test on independent days.

#### Cued-fear conditioning

Experiments were performed at the same time of day during the light phase using 3–4-month-old male and female mice (n = 14–19 mice/genotype, with balanced representation of both sexes), following previously described procedures (Fernández de Sevilla et al., 2022). Training and retrieval sessions were video-recorded. Mice were placed in a shuttle box (AccuScan Instruments) and allowed to habituate for 120 s before exposure to an auditory neutral stimulus (CS+; a tone of 80 dB for 30 s, presented together with an aversive unconditioned stimulus (US; electrical foot shock, 0.3 mA, 2 s). The foot shock was delivered during the final 2 s of the auditory stimulus, so that both stimuli co-terminated. In experiments employing the PTSD-inducing protocol, shock intensity was increased to 2.5 mA (Siegmund and Wotjak, 2007). Fear acquisition consisted of five CS–US pairings separated by 90-s intertrial intervals (CS−). Following the final pairing, animals remained in the conditioning chamber for an additional 5 min before being returned to their home cages.

Twenty-four hours later, fear conditioning was tested placing the animals in a different context using the same protocol but without the unconditioned stimulus (US). As a consequence of the US–CS association, mice displayed freezing behavior, which was scored for every trial (CS+), and intertrial (CS-). Contextual cues, including chamber wall configuration and illumination, were changed between training and retrieval sessions, and the apparatus was cleaned between sessions.

Freezing behavior was defined as the absence of movement except for respiration and heartbeat. Freezing was quantified through continuous visual inspection of video recordings by experimenters blinded to group allocation. Freezing duration was assessed separately during each CS+ presentation and the corresponding intertrial intervals (CS−), according to the predefined observation periods, and expressed as the percentage of time spent freezing within each interval.

For fasting experiments, food was withdrawn immediately after fear-conditioning acquisition, and mice remained food-deprived for 24 h, with water available ad libitum. Conditioned fear was subsequently assessed using the retrieval protocol described above. Fed control animals retained unrestricted access to standard chow.

#### Open field test

Exploratory behavior and locomotor activity were assessed using a CIBERTEC Actimeter system (Actimeter XYZ-8; CIBERTEC S.A., Spain). Mice were individually placed in an open-field arena (42 × 42 × 30 cm) for 5 min. Horizontal and vertical activity were recorded using infrared photocell arrays and analyzed with the MUX_XYZ16L-8 software. Time spent in specific areas of the arena, including the central region, was quantified to assess exploratory behavior, as previously described (Zegarra-Valdivia et al., 2019).

#### Elevated plus maze

Anxiety-like behavior was assessed using an elevated plus maze (Med Associates, USA) positioned 40 cm above the floor. The apparatus consisted of four arms arranged in a cross-shaped configuration: two opposing closed arms (30 cm long × 5 cm wide × 15.25 cm high) enclosed by protective walls, and two opposing open arms (30 cm long × 5 cm wide) without protective walls. Each mouse was individually placed in the central platform and allowed to explore the maze freely for 5 min. Time spent in the open and closed arms was recorded using the Video Tracking Plus Maze Mouse system (Med Associates, USA). Time spent in the open arms was used as an indicator of anxiety-like behavior, with reduced open-arm exploration reflecting increased anxiety-like responses. Behavioral data were analyzed as previously described (Munive et al., 2019).

### Optogenetic stimulation of orexin neurons

Orexin neurons were optogenetically activated using an AAV8-HCRT-ChR2-EGFP vector injected into the orexin-containing region of the lateral hypothalamus, at the stereotaxic coordinates described above (AP, −1.4 mm; ML, ±0.9 mm; DV, −5.4 mm). Animals underwent behavioral testing four weeks after viral delivery.

Twenty-four hours after fear conditioning, mice were tested in a modified context. Following a 2-min habituation period, freezing behavior was recorded during a 3-min light-off period followed by a 3-min light-on period. Blue-light stimulation was delivered at 470 nm through an implanted optical fiber, using 20-Hz stimulation with 35-ms pulses. Freezing behavior was quantified from video recordings and compared between stimulation conditions and experimental groups.

### Quantitative PCR

Total RNA isolation from brain tissue was carried out with Trizol. One mg of RNA was reverse transcribed using High Capacity cDNA Reverse Transcription Kit (ThermoFisher Scientific, Waltham, MA, USA) according to the Manufacturer’s instructions. For quantification of specific genes, total RNA was isolated and transcribed as above and 62.5 ng of cDNA was amplified using TaqMan probes for mouse IGF-1receptor (IGF-1R) and Insulin receptor (InsR), and rRNA 18S as endogenous control (ThermoFisher Scientific). Each sample was run in triplicate in 20 µL of reaction volume using TaqMan Universal PCR Master Mix according to the manufacturer’s instructions (ThermoFisher Scientific). All reactions were performed in a 7500 Real Time PCR system (ThermoFisher Scientific). Quantitative real time PCR analysis was carried out as described^8^ – Indicar cita. Results were expressed as relative expression ratios on the basis of group means for target transcripts versus reference 18S transcript. At least five independent experiments were done.

### Immunocytochemistry

Immunocytochemistry was performed as described in detail before (Fernández de Sevilla et al., 2022). Primary antibody was omitted as control. Mice were perfused trans-cardially under deep anesthesia (sodium pentobarbital, 50 mg/kg, intraperitoneously) with 100 ml of saline buffer 0.9% followed by 100 ml of 4% paraformaldehyde (PFA) in 0.1N, pH 7.4 phosphate buffer (PB), 90 min after the cued-fear conditioning test, the recall context, or under basal conditions. This time point was selected as optimal to see c-Fos staining. Brains were removed, post-fixed overnight at 4°C in the same fixative, and cut at 50 µm thick sections on a vibratome (Leica VT 1000S). Sections were kept at 4°C, immersed in 0.1N PB with 0.02% sodium azide, until processing. Serial coronal free-floating sections were rinsed in 0.1N PB for 10 min, and a blocking solution containing 10% normal donkey serum, and 0.4% Triton X-100 in 0.1N PB (PB-T) was added and maintained at room temperature for 2 h.

Thereafter, sections were incubated overnight at 4°C in the same solution with the corresponding primary antibodies. The next day, sections were washed 3 times with PB-T, and incubated with secondary antibodies for 2 h at room temperature. After the incubation, slices were washed 3 times with PB-T and incubated 5 min with Hoechst (1:500 dilution, Life Technologies). Finally, sections were washed 3 times with PB and mounted onto glass slides coated with gelatin in Gelvatol mounting medium. Images were taken with confocal microscopy (SP5, Leica Microsystems, Germany).

### Image analysis

Analyses were carried out with Imaris software (Bitplane), which allows to examine images in 3D. With the images taken at 20x magnification, we created spots with an estimated XY size of 15 µm to count orexin^+^ or TH^+^ neurons in their channel (red), and next we added a filter corresponding to the c-Fos channel (green) to calculate the number of orexin^+^/c-Fos^+^ and TH^+^/c-Fos^+^ cells.

### RNAscope

RNAscope (2.5 HD Detection kit—Red; #322350; ACD, USA) with an IR-specific probe combined with immunocytochemistry with anti-GFAP antibody was performed to confirm reduced IR expression in GFAP+ cell 1 month after virus injection in the tail (Suppl Fig 1C,D).

### Statistics

Statistical analyses were performed using GraphPad Prism 10.4.1 (GraphPad Software; RRID). Data are presented as mean ± SEM unless otherwise indicated. Normality was assessed using the Shapiro–Wilk test. Comparisons between two groups were performed using paired or unpaired Student’s t-tests, Welch’s t-test, or the Mann– Whitney U test, as appropriate. Comparisons involving multiple groups or experimental factors were performed using one-, two-, or three-way ANOVA, including repeated-measures designs, or mixed-effects models fitted by restricted maximum likelihood (REML), followed by appropriate post hoc tests. All statistical tests were two-sided, with significance set at p < 0.05. Sample sizes and specific statistical procedures are indicated in the corresponding figure legends.

## Acknowledgements

We are thankful to M. Garcia for technical support. This work was funded by a grant from Ciberned, and from grant PID2019-104376RB-I00 (AEI/FEDER; MICINN).

JAZ-V is an IKUR fellow and MEFdS was supported by fellowship to grant SAF2016-76462 (AEI/FEDER; MICINN)

## Authoŕs contributions

JAZV and MEFdS performed and designed experiments, analyzed results, and wrote parts of the manuscript. JP performed and designed experiments, analyzed results, and wrote parts of the manuscript. ZK performed experiments. ITA designed the study, analyzed and interpreted results, and wrote the manuscript.

## Declaration of Generative IA

During the preparation of this manuscript, generative AI and AI-assisted technologies (ChatGPT, GPT-6; OpenAI) were used to improve grammatical accuracy, clarity, readability, and scientific English, as well as to assist with the graphical refinement and visual harmonization of author-designed experimental schematics and conceptual illustrations. All AI-assisted outputs were critically reviewed and revised by the authors to ensure accuracy and consistency with the original scientific content. The authors take full responsibility for the content of the final manuscript.

## LEGENDS TO FIGURES

**Supplementary Figure 1.** Food and drink consumption, astrocytic insulin receptor expression, and the effects of fasting on conditioned fear. **A**, Wild-type mice exposed to palatable food consumed significantly more food than mice exposed to regular food during a 2-h period (Regular food, n = 20; palatable food, n = 24; two-tailed Mann– Whitney U test, U = 114, p = 0.0012). **B,** Wild-type mice exposed to a palatable drink consumed significantly greater volumes than mice receiving a regular drink during a 2-h period (Regular drink, n = 15; palatable drink, n = 25; two-tailed Mann–Whitney U test, U = 15, p < 0.0001). **C,** Combined GFAP immunostaining (green) and insulin receptor (IR) mRNA detection by RNAscope in situ hybridization (red) in IRfl/fl (BIR) control mice and BIR-GFAP-Cre mice following systemic administration of an AAV-Cre vector. Cell nuclei were counterstained with DAPI (blue). Enlarged images correspond to regions indicated by white squares. Scale bars: 25 µm and 5 µm (insets). **D,** IR expression was significantly decreased in BIR-GFAP-Cre mice compared with BIR controls, as determined by quantification of IR RNAscope puncta in GFAP-positive cells (BIR, n = 4; BIR-GFAP-Cre, n = 4; unpaired two-tailed Student’s t-test, t(6) = 5.102, p = 0.0022). **E,** Food deprivation for 24 h increased conditioned freezing during CS+ presentations in wild-type mice compared with fed controls (Fed, n = 8; Fasted, n = 8; CS− measurements available for n = 7 per group). A repeated-measures mixed-effects model (REML) revealed a significant stimulus period × feeding condition interaction (F(1,12) = 22.44, p = 0.0005). Šidák-adjusted comparisons confirmed increased freezing during CS+ (p < 0.0001), whereas the difference during CS− was not significant (p = 0.1517). Data are presented as mean ± SEM. **p < 0.01, ****p < 0.0001.

**Supplementary Figure 2.** Astrocytic insulin receptor knockdown increases c-Fos expression in fear-related nuclei following fear conditioning. **A**, Representative micrographs showing double immunostaining for c-Fos (green) and orexin (red) in the lateral hypothalamus (upper panels), and c-Fos (green) and tyrosine hydroxylase (TH; red) in the locus coeruleus (LC; lower panels) of BIR Control mice (left) and BIR-GFAP-Cre mice (right) following fear conditioning. **B,** The percentage of orexin-positive neurons coexpressing c-Fos was significantly increased in BIR-GFAP-Cre mice compared with BIR controls (BIR Control, n = 4; BIR-GFAP-Cre, n = 4; unpaired two-tailed Student’s t-test, t(6) = 3.762, p = 0.0094). **C,** The percentage of TH-positive neurons coexpressing c-Fos in the LC was significantly increased in BIR-GFAP-Cre mice compared with BIR controls (BIR Control, n = 4; BIR-GFAP-Cre, n = 4; unpaired two-tailed Student’s t-test, t(6) = 2.558, p = 0.0430). Data are presented as mean ± SEM, with individual points representing mice. Scale bars: 100 µm. *p < 0.05, **p < 0.01.

**Supplementary Figure 3.** Hippocampal c-Fos expression and chemogenetic inhibition of lateral hypothalamic astrocytes. **A**, No significant differences in c-Fos-positive cell density were observed across the hippocampal regions examined (CA1, CA2, CA3, CA4, and dentate gyrus) among the experimental groups (n = 3 mice per group). CA, cornu ammonis; DG, dentate gyrus. **B,** Chemogenetic inhibition of lateral hypothalamic (LH) astrocytes using inhibitory DREADDs and clozapine-N-oxide (CNO) administration in fed wild-type mice did not significantly alter conditioned freezing responses compared with vehicle-treated controls (n = 5–7 mice per group). Data are presented as mean ± SEM.

