## Supplementary figures and images for "Insulin Modulates Food Valence Through Astrocytes"

### Supplementary Figures

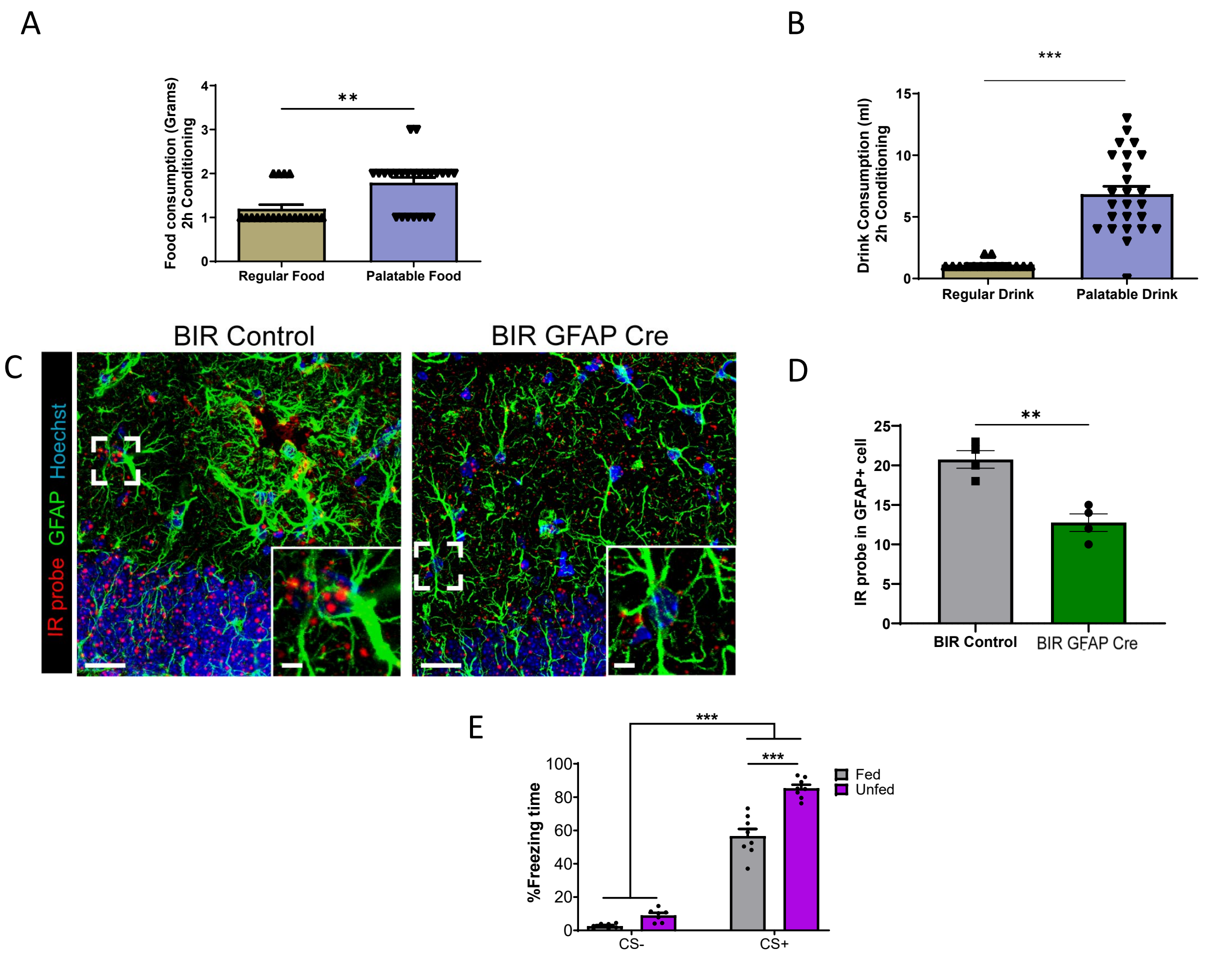


Supplemmentary Figure 1

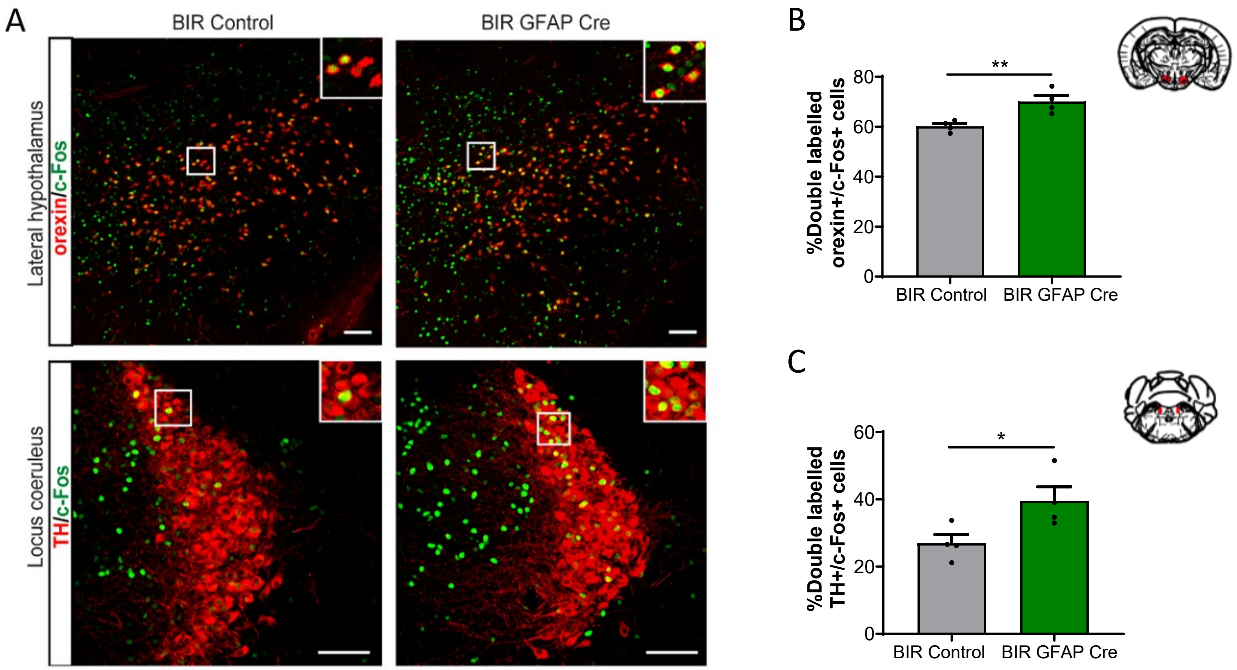


Supplemmentary Figure 2

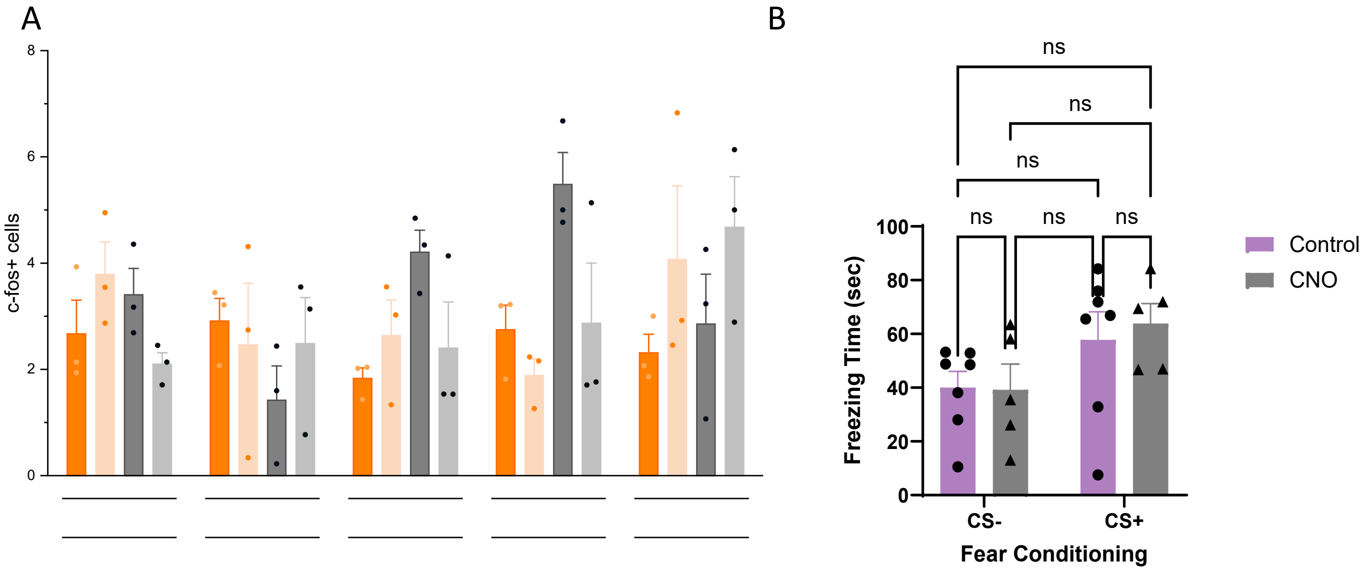


Supplemmentary Figure 3
